# Time-gated detection of protein-protein interactions with transcriptional readout

**DOI:** 10.1101/166462

**Authors:** Min Woo Kim, Wenjing Wang, Mateo I. Sanchez, Robert Coukos, Mark Von Zastrow, Alice Y. Ting

## Abstract

Transcriptional assays such as yeast two hybrid, split ubiquitin, and Tango that convert transient protein-protein interactions (PPIs) in cells into stable expression of transgenes are powerful tools for PPI discovery, high-throughput screens, and analysis of large cell populations. However, these assays frequently suffer from high background and they lose all information about PPI dynamics. To address these limitations, we developed a light-gated transcriptional assay for PPI detection called PPI-FLARE (PPI-Fast Light- and Activity-Regulated Expression). PPI-FLARE requires *both* a PPI to deliver TEV protease proximal to its cleavage peptide, *and* externally-applied blue light to uncage the cleavage peptide, in order to release a membrane-tethered transcription factor (TF) for translocation to the nucleus. We used PPI-FLARE to detect the ligand-induced association of 12 different PPIs in living mammalian cells, with a temporal resolution of 5 minutes and a ±ligand signal ratio up to 37. By systematically shifting the light irradiation window, we could reconstruct PPI time-courses, distinguishing between GPCRs that engage in transient versus sustained interactions with the cytosolic effector arrestin. When combined with FACS, PPI-FLARE enabled >100-fold enrichment of cells experiencing a specific GPCR-arrestin PPI during a short 10-minute light window over cells missing that PPI during the same time window. Due to its high specificity, sensitivity, and generality, PPI-FLARE should be a broadly useful tool for PPI analysis and discovery.

---

Protein-protein interactions (PPIs) are central to cellular signal transduction. Consequently, many assays have been developed to detect and study them, particularly in the context of living cells, where native PPIs are unperturbed by cell lysis, detergents, fixatives, or dilution. For example, FRET^1^, BRET^2^, fluorescence correlation spectroscopy (FCS)^3^, protein complementation assays (PCAs)^4^, and fluorescence relocalization assays^5,6^ have all been used to visualize PPI dynamics in living cells. Though information-rich, these assays have the downside of being labor-intensive, requiring high-content or time-lapse microscopy, and are consequently difficult to perform on a large scale. This makes them non-optimal for PPI discovery, for adaptation to high-throughput screens, or for analysis of large cell populations such as in complex tissue. Instead, real-time assays are more suited for the focused study of a small number of known PPIs under a small set of conditions or in a small number of cells.

A separate class of assays detects PPIs by signal integration rather than real-time imaging, and produces gene transcription as the readout. Examples include the yeast two hybrid assay^7^, the split ubiquitin assay^8^, and Tango^9–11^. Benefits of these assays include scalability (because real-time microscopy is not needed), versatility of read out (transcription of a fluorescent protein or an antibiotic resistance gene, for example), and high sensitivity due to signal amplification. These properties have led to the widespread use of integrative PPI assays for PPI discovery and drug screening. However, these assays have two major drawbacks. First, because they integrate PPI events over long time periods, typically 18 hours to days (i.e., over the entire time that the tools are expressed in cells), they often produce high background, leading to high rates of false discovery. Second, these assays lose information about PPI dynamics. It is impossible to know, for instance, whether a PPI event occurred during a specific 5-minute time window of interest or at another time.

To address these limitations, and greatly expand the potential utility and versatility of PPI assays that work by signal integration and transcriptional readout, we report here a new tool, called PPI-FLARE, for “Protein-Protein Interactions detected by Fast Light and Activity-Regulated Expression”. PPI-FLARE detects interactions in living cells between specific protein pairs of interest (proteins A and B in Fig. 1A), and produces gene transcription as a result. However, in contrast to previous tools, PPI-FLARE is also gated by externally-applied blue light. Hence, transcriptional activation requires both protein A-protein B interaction *and* light, the latter of which can be applied during any user-specified time window. This generic and non-invasive form of temporal gating enables PPI-FLARE to re-capture PPI dynamics to some extent, and reduces background signal overall, while preserving the tremendous benefits of transcriptional readout.

**Figure 1:**
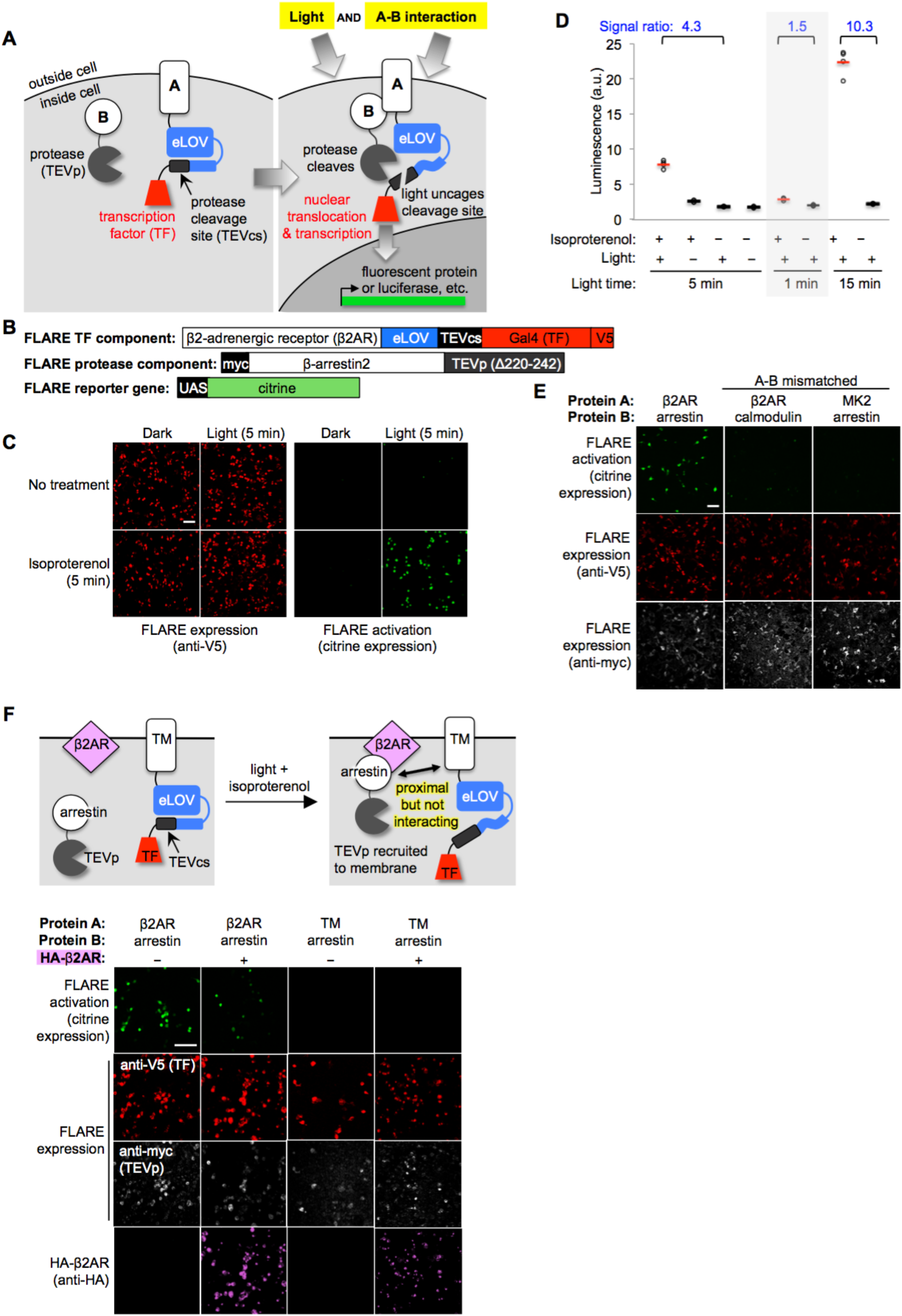
Design of PPI-FLARE and application to light- and agonist-dependent detection of β2-adrenergic receptor (β2AR)-β-arrestin2 interaction. (**A**) Scheme. A and B are proteins that interact under certain conditions. Protein A is membrane-associated and is fused to a light-sensitive eLOV domain^12^, a protease cleavage site (TEVcs), and a transcription factor (TF). These comprise the “FLARE TF component.” Protein B is fused to a truncated variant of TEV protease (TEVp) (“FLARE protease component”). When A and B interact (right), TEVp is recruited to the vicinity of TEVcs. When blue light is applied to the cells, eLOV reversibly unblocks TEVcs. Hence, the coincidence of light *and* A-B interaction permits cleavage of TEVcs by TEVp, resulting in the release of the TF, which translocates to the nucleus and drives transcription of a reporter gene of interest. (**B**) PPI-FLARE constructs for studying the β2AR-β-arrestin2 interaction. V5 and myc are epitope tags. UAS is a promoter recognized by the TF Gal4. (**C**) Imaging of FLARE activation by β2AR-β-arrestin2 interaction under four conditions. HEK 293T cells were transiently transfected with the three FLARE components shown in (B). β2AR-β-arrestin2 interaction was induced with addition of 10 µM isoproterenol for 5 minutes. Light stimulation was via 467 nm LED at 60 mW/cm^2^ and 10% duty cycle (0.5 second of light every 5 seconds) for 5 minutes. Nine hours after stimulation, cells were fixed and imaged. (**D**) Same as (C), but HEK 293T cells were stably expressing the FLARE protease component and transiently expressing FLARE TF component and UAS-luciferase. Results of shorter and longer irradiation times are also shown. ± isoproterenol signal ratio was quantified for each time point. Each datapoint reflects one well of a 96-well plate containing >6,000 transfected cells. Four replicates per condition. (**E**) FLARE is specific for PPIs over non-interacting protein pairs. Same experiment as in (C), except arrestin was replaced by calmodulin protein (which does not interact with β2AR) in the second column, and β2AR was replaced by the calmodulin effector peptide MK2 (which does not interact with arrestin) in the third column. Anti-myc and anti-V5 antibodies stain for the FLARE protease and TF components, respectively. (**F**) FLARE is activated by direct interactions and not merely proximity. Top: experimental scheme. To drive proximity but not interaction, we created FLARE constructs in which A and B domains were a transmembrane (TM) segment of the CD4 protein, and β-arrestin2, respectively. TM and arrestin do not interact. HEK 293T cells expressing these FLARE constructs were also transfected with an expression plasmid for HA-tagged β2AR. Upon isoproterenol addition, β-arrestin2-TEVp is recruited to the plasma membrane via interaction with β2AR, but it does not interact directly with the FLARE TF component. Bottom: Images of HEK 293T cells 9 hours after stimulation with isoproterenol and light (for 5 minutes). The last column shows the experiment depicted in the scheme. The first two columns are positive controls with FLARE constructs containing β2AR and β-arrestin2 (which do interact). The third column is a negative control with omission of the HA-β2AR construct. Anti-V5, anti-myc, and anti-HA antibodies stain for FLARE TF component, FLARE protease component, and HA-β2AR proteins, respectively. All scale bars, 100 µm.

Here we describe the development of PPI-FLARE, its characterization in living mammalian cells, and its application to a range of PPIs, including eight different GPCRs. We use PPI-FLARE to compare the temporal profiles of different GPCRs interacting with β-arrestin2 in response to various ligands. Finally, we show that PPI-FLARE can enable FACS-based enrichment of cells that experienced specific transient (10-minute long) PPI events >9 hours earlier. This lays the foundation for the use of PPI-FLARE for genetic screens and PPI discovery on a genome-wide scale.

## Design and optimization of PPI-FLARE

To design PPI-FLARE, we built upon our recently reported Ca-FLARE tool, which is a transcription factor gated by the coincidence of light and high calcium^12^. As shown in Figs. 1A-B, PPI-FLARE has three components: a transcription factor (TF) that is tethered to potential protein interaction partner “A” as well as to the plasma membrane (or other intracellular membrane, sequestered from the nucleus) via a protease cleavage site (“TEVcs” for tobacco etch virus cleavage site); a protease (“TEVp” for tobacco etch virus protease) that is fused to potential protein interaction partner “B”; and a reporter gene whose transcription is triggered by TF translocation to the nucleus. TF release from the plasma membrane via TEVcs cleavage requires the coincidence of two events: protein A-protein B interaction *and* externally-applied blue light. This is because, first, the TEVp protease is tuned to have low affinity for its TEVcs substrate and cleavage is only significant when the two are brought into proximity via A-B interaction. Second, the TEVcs is caged by an “evolved LOV domain” (eLOV^12^), which we previously generated by directed evolution to tightly cage the TEVcs sequence and prevent its cleavage in the dark state. A brief (<1 sec) pulse of 467 nm light, however, alters the eLOV protein conformation, unblocking the adjacent TEVcs and allowing cleavage. After ~100 seconds without blue light, eLOV spontaneously reverts to its dark state conformation and again blocks TEVcs.

To test PPI-FLARE with a well-established cellular PPI, we selected the β2-adrenergic receptor (β2AR), a 7-transmembrane GPCR that recruits the soluble cytosolic effector protein β-arrestin2 after stimulation with small-molecule ligands such as isoproterenol. We cloned β2AR and β-arrestin2 genes in the protein A and protein B positions of PPI-FLARE, respectively (Fig. 1B). Note that β2AR is not fused to the tail domain of the vasopressin receptor^9,13^ or any other motif that artificially boosts its recruitment of arrestin; only native genes are utilized. HEK 293T cells expressing these FLARE components were stimulated for 5 minutes with both 467 nm light (50 mW/cm^2^) and isoproterenol. Under these conditions, we expect FLARE components to interact and release the transcription factor, Gal4, from the plasma membrane. Consequently, we should observe transcription and translation of the Citrine fluorescent protein reporter gene. Nine hours after stimulation of the HEK cells, we fixed them and imaged Citrine expression. Fig. 1C shows robust Citrine expression in the light- and isoproterenol-treated cells. Essential for a functional “AND” gate, we detected negligible Citrine expression in the light-only or isoproterenol-only cells.

Because PPI-FLARE is a transcriptional tool, the readout can be any gene of our choosing. Luciferase is frequently the readout of choice for high-throughput assays, because it is easy to detect and quantify. By repeating the above experiment, but replacing the UAS-Citrine gene with a UAS-luciferase gene, and quantifying luminescence on a platereader, we again detected a robust increase in reporter gene transcription in light- and isoproterenol-treated cells compared to light-only or isoproterenol-only cells (Fig. 1D).

In an effort to further optimize PPI-FLARE, we explored alternative TEVcs sequences and alternative LOV domain sequences. Alternative TEVcs sequences, particularly with changes at the P1´ position immediately following the cleavage site, have the potential to alter both k_cat_ and K_m_ of the TEVp-TEVcs interaction. For optimal PPI-FLARE performance, we favor high k_cat_, for high signal, and high K_m_, for low background and strong proximity-dependence of the cleavage. Hence, we explored TEVcs variants with P1´ = M, Q or Y because these have previously been shown to tune TEVp-TEVcs kinetics^12,14^.

Supplementary Fig. 1 shows that P1´ = M displayed the highest signal and strongest proximity dependence in the context of β2AR-β-arrestin2 FLARE, and hence we selected this TEVcs sequence for all subsequent experiments in this study.

While our evolved eLOV provides robust light caging of TEVcs, with light/dark signal ratios ranging from 40 to 120 in HEK 293T cells and cultured neurons^12^, a new LOV sequence with improved properties over the original LOV (which served as the template for our eLOV evolution) has been reported^13,15^. We thus performed a side-by-side comparison of this new LOV domain (called “iLID”) to our eLOV, in the context of β2AR-β-arrestin2 PPI-FLARE. In addition, we constructed two different hybrid LOV domains (hLOV1 and hLOV2) that combine features of eLOV and iLID into a single gene (Supplementary Fig. 2A). Supplementary Fig. 2 shows that with short stimulation times, or low FLARE component expression levels, eLOV is the best light gate, because it gives the highest signal in the +light condition (Supplementary Fig. 2B and 2D). When FLARE components are highly expressed, and the stimulation time is increased from 5 minutes to 15 minutes, however, hLOV1 is the best light gate, because it gives the lowest signal in the no-light condition (i.e., best steric protection of TEVcs in the dark state, Supplementary Fig. 2C). Because our study requires short tagging time periods, we selected eLOV for all subsequent experiments in this study. For applications that require minimization of dark state leak, however, hLOV1 is worth considering as an alternate light gate.

## Further characterization of PPI-FLARE in mammalian cells

We compared different light stimulation times and found that 5 minutes is sufficient to give a 4.3-fold increase in FLARE-driven luciferase expression (Fig. 1D). 1 minute of light is too short, and one can optionally increase the stimulation time to 15 minutes for further boost the ±isoproterenol signal ratio to 10.3. To test FLARE-PPI specificity, we replaced one protein partner in the β2AR-β-arrestin2 FLARE pair with a non-interacting protein. Fig. 1E shows that the mismatched pairs β2AR-calmodulin and MK2-β-arrestin2 fail to drive Citrine expression.

We wondered whether mere recruitment of TEVp to the vicinity of TEVcs, without direct complexation mediated by the A-B PPI, would be sufficient to give proteolytic cleavage and TF release. To test this, we generated a mismatched FLARE pair using β-arrestin2 (protein B) and the transmembrane domain of CD4 (protein A). We transfected HEK cells with these FLARE constructs along with a V5-tagged β2AR expression plasmid. We then drove arrestin-TEVp translocation from the cytosol to the plasma membrane by addition of isoproterenol (Supplementary Fig. 3A). Fig. 1F and Supplementary Fig. 3B show that, even when arrestin-TEVp is recruited to the immediate vicinity of the TM (CD4) FLARE TF component, no FLARE activation (citrine expression) is observed. This demonstrates that FLARE is a very specific detector of direct A-B *interaction*, and is not activated merely by A-B *proximity*.

How efficient is PPI-FLARE? How much TEVcs cleavage occurs after our standard stimulation time of 5 minutes? To quantify this, we used the β2AR-β-arrestin2 FLARE constructs in Fig. 1B, and lysed the cells immediately after stimulation to examine the FLARE proteins by Western blot. Supplementary Fig. 4 shows that 30% of total TEVcs is cleaved after 5 minutes of light and isoproterenol, and the cleavage yield increases to 48% after 30 minutes of stimulation.

For most of our experiments, we used a blue LED array to deliver light to cells. However, many laboratories do not have access to such a light source. The light power required to uncage the LOV domain is very weak, <0.5 mW/cm^2^ [ref. 16]. Thus we tested if PPI-FLARE could be stimulated just as well by ordinary room light. Supplementary Fig. 5 shows that the signal ratios obtained are the same whether the light source is a blue LED or ambient room light. In some experiments in our study, we also used a broad-wavelength “daylight lamp” to deliver light to FLARE-expressing cells. When we did not wish for light exposure, we kept the cells covered in aluminum foil and worked in a dark room with a red light source.

## PPI-FLARE can be generalized to a variety of PPIs

To test the generality of PPI-FLARE, we replaced β2AR and arrestin with a variety of other PPI pairs (Fig. 2A). These include two different GPCRs - dopamine receptor D1 (DRD1) and neuromedin B receptor (NMBR) - that also recruit arrestin; a tyrosine kinase receptor that recruits Grb2 protein upon stimulation with a growth factor; the FKBP-FRB protein pair whose interaction is induced by rapamycin; and the light-regulated CRY2-CIBN protein pair. In contrast to β2AR, the latter four proteins are all soluble proteins that cannot by themselves sequester the TF component of FLARE from the nucleus. To ensure that the TF remains in the cytosol in the basal state, we fused transmembrane domains to both FKBP and CIBN.

**Figure 2:**
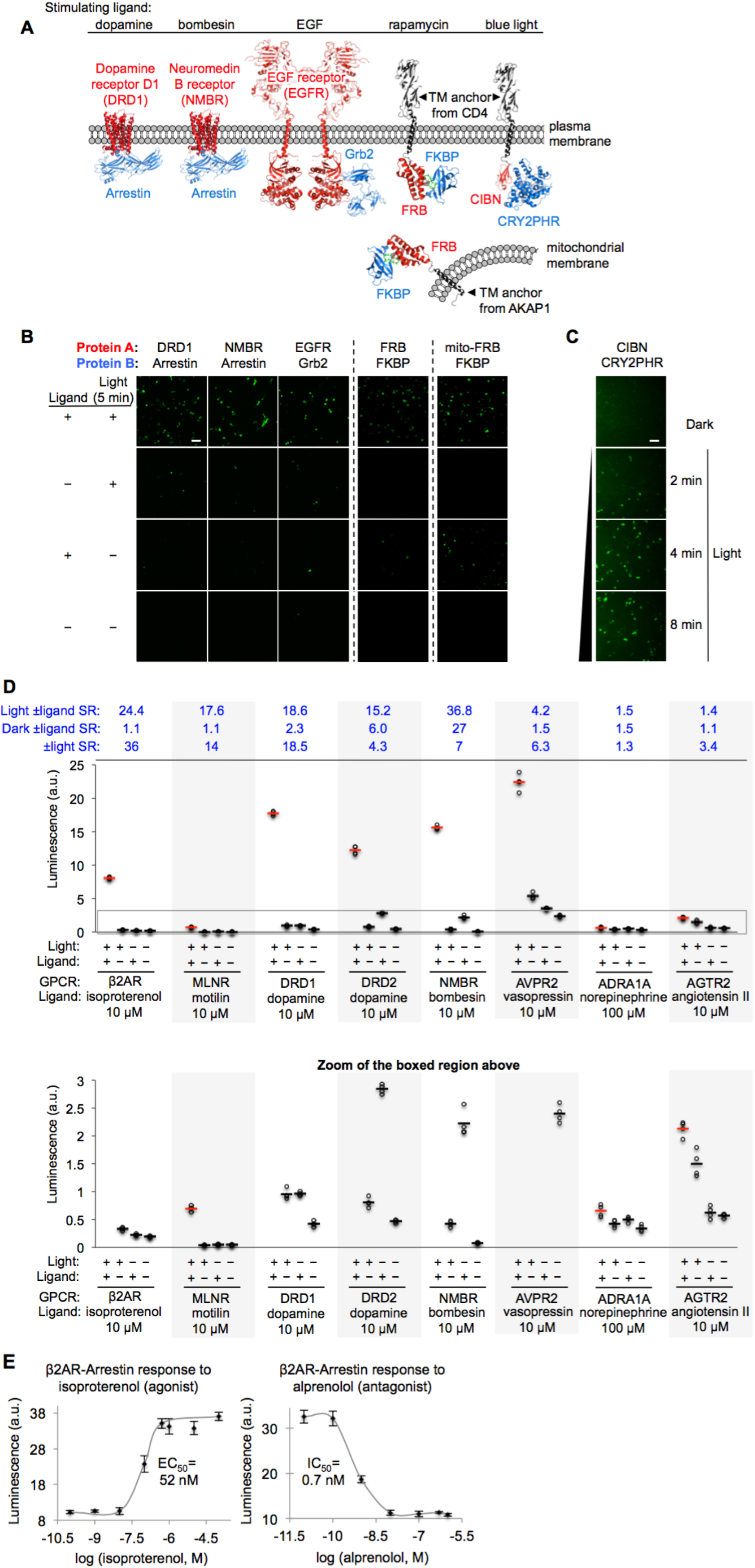
PPI-FLARE can be applied to a variety of PPIs. (**A**) PPI pairs studied with FLARE. DRD1 and NMBR are GPCRs that interact with β-arrestin2. EGFR is a receptor tyrosine kinase that recruits Grb2 upon stimulation with EGF ligand. FKBP and FRB are soluble proteins that heterodimerize upon addition of the drug rapamycin; to keep FRB FLARE out of the nucleus in the basal state, we fuse it to either a plasma membrane anchor (TM from CD4) or a mitochondrial membrane anchor (TM from AKAP1). CIBN-CRY2PHR is a light-inducible PPI^40^. (**B**) FLARE data corresponding to PPIs depicted in (A). FLARE constructs were the same as those shown in Fig. 1B, except β2AR and β-arrestin2 were replaced by the A and B proteins indicated, respectively. HEK 293T cells transiently expressing FLARE constructs were stimulated with light and the ligand indicated in (A) for 5 minutes, then fixed and imaged 9 hours later. Citrine fluorescence images are shown. Dashed lines separate experiments that were performed separately and shown with different Citrine intensity scales. Scale bar, 100 µm. (**C**) FLARE detection of CIBN-CRY2PHR interaction. Blue light (467 nm, 60 mW/cm^2^, 33% duty cycle (2 seconds light every 6 seconds)) simultaneously uncages the eLOV domain and induces the CIBN-CRY2PHR interaction. Scale bar, 100 µm. (**D**) FLARE applied to 8 different GPCRs. HEK 293T cells were prepared as in Fig. 1D. The FLARE protease component is β-arrestin2-TEVp. The FLARE TF component contains the indicated GPCR (no vasopressin V2 domain). Light (ambient) and ligand were applied for 15 minutes total, then cells were analyzed for luciferase activity 9 hours later. Four replicates per condition. ± ligand signal ratios (SR) and ± light signal ratios for each GPCR quantified across top. The boxed region is enlarged in the bottom. (**E**) Isoproterenol and alprenolol dose-response curves with β2AR-β-arrestin2 FLARE readout. HEK 293T cells were prepared and stimulated as in Fig. 1D, with 5 minute light window. Four replicates per concentration. Errors, STD. EC_50_ of 52 nM for isoproterenol and IC_50_ of 0.7 nM for alprenolol are close to published values^41,42^.

Figs. 2B-C show that all protein pairs tested gave clear light- and ligand-dependent gene expression. Some background signal was observed in the no-light/+rapamycin condition for FRB/FKBP, but this is to be expected given that rapamycin dissociates very slowly (t_1/2_ ~17.5 hours [ref. 17]) and consequently, the TEVp/TEVcs domains of these FLARE constructs remain in close proximity for the entire 9 hr transcription/translation window. To test whether the TF component of FLARE would also work in a different subcellular region besides the plasma membrane, we created a FKBP FLARE construct localized to the outer mitochondrial membrane (OMM) (via fusion to the N-terminal 53-amino acid OMM targeting sequence of AKAP1). Fig. 2B shows that this FLARE pair gives rapamycin- and light-dependent gene expression as well.

GPCRs are a protein class of special interest due to their central role in signal transduction, their rich pharmacology, and their prevalence as therapeutic targets. Though ligand-activated GPCRs differ in their downstream recruitment of various G proteins (G_i_, G_s_, or G_q_), nearly all of them recruit the effector arrestin as part of their desensitization pathway^18^. As a consequence, assays that read out arrestin recruitment to GPCRs can generically detect activation of a wide range of GPCRs. We were interested to know whether PPI-FLARE could detect the ligand-dependent recruitment of arrestin to a range of GPCRs. In Fig. 2D, we selected eight different GCPRs and tested them in FLARE under four conditions. The first six GPCRs gave robust ligand-dependent luciferase expression, five of them with ±ligand signal ratios >15. Such large signal ratios are especially encouraging given that previous work has shown that some of these GPCRs (e.g., DRD2, NMBR and AVPR2) have significant interactions with arrestin prior to ligand addition^11^, suggesting that background signal (FLARE-driven luciferase expression in the no ligand condition) might be a concern. Despite this, the high sensitivity and specificity of FLARE permitted facile differentiation between the +ligand and no ligand states.

The last two GPCRs we surveyed, AGTR2 and ADRA1A, are not known to recruit arrestin, and previous assays (e.g., PRESTO-Tango^11^) failed to produce signal. Our FLARE ±ligand signal ratios were small for these GPCRs (1.5 and 1.4, respectively, Fig. 2D) but statistically significant (p <0.001 for both). Thus FLARE enables us to conclude that these receptors do indeed recruit arrestin, but to a much lesser extent than other GCPRs.

Two GPCRs in our panel, DRD2 and NMBR, exhibited higher background than others in the +ligand/no light state (leading to higher ±ligand (dark state) signal ratios in Fig. 2B), similar to the FKBP-FRB pair in Fig. 2B. We hypothesize that this is because these two GPCRs interact with arrestin in a sustained rather than transient manner after ligand addition, similar to how rapamycin produces sustained FKBP-FRB interaction. When a ligand causes A-B partners in FLARE to engage for hours, some TEVp-catalyzed cleavage of TEVcs will occur, even in the absence of light (eLOV protection of TEVcs is not perfect), leading to background transcription in the +ligand/no light state. GPCRs known to recruit arrestin transiently, such as β2AR, do not exhibit this phenomenon (Fig. 2B). Interestingly, our time course study on DRD2 and NMBR below (Fig. 3E) is consistent with this interpretation.

**Figure 3:**
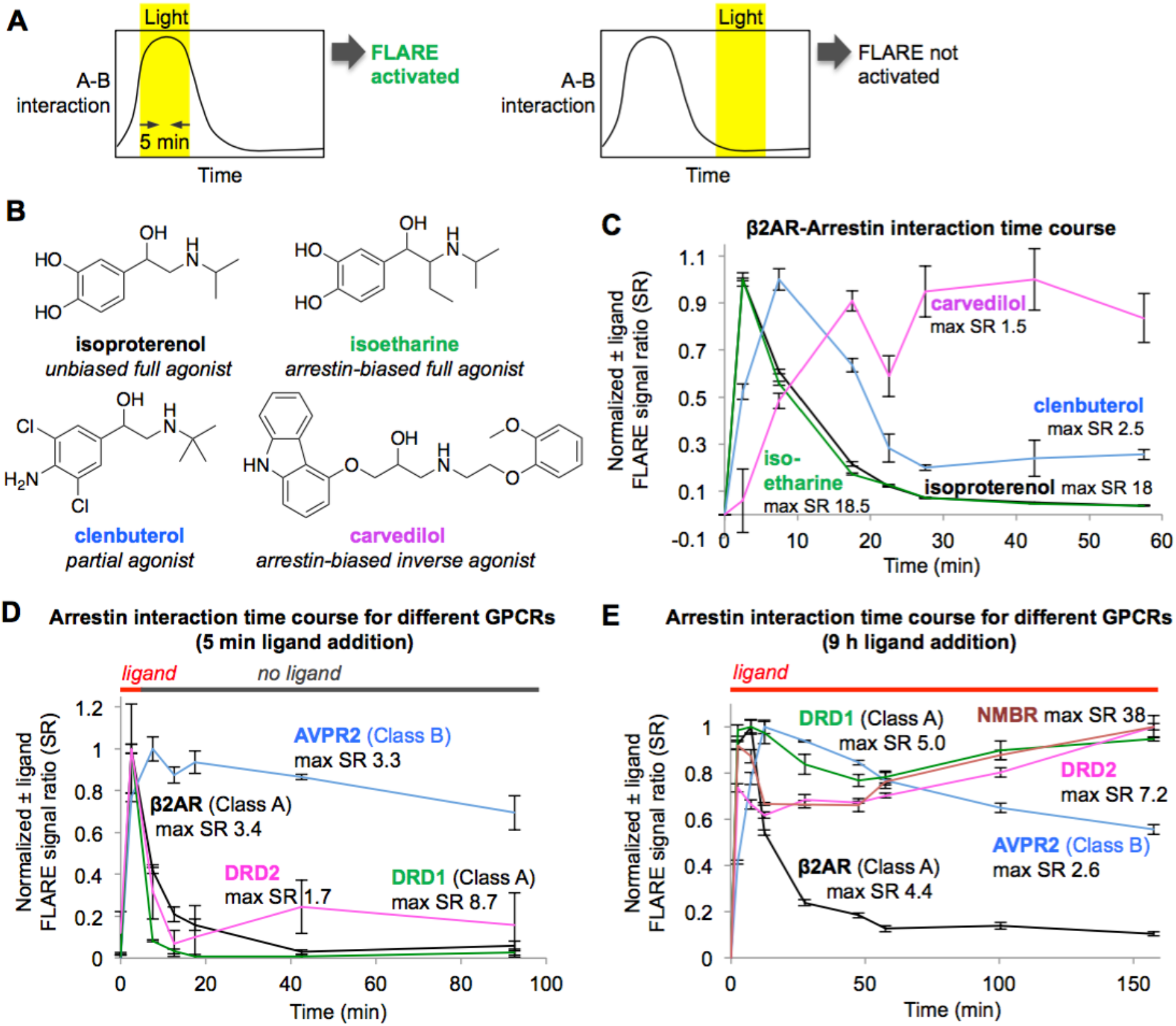
Light gating of PPI-FLARE permits analysis of the dynamic GPCR-β-arrestin2 interaction. (**A**) Scheme. By shifting the light window, it is possible to read out different time regimes of protein A-protein B interaction. On the left, light coincides with a period of high A-B interaction, resulting in FLARE activation and transcription of a reporter gene. On the right, light coincides with a period of low A-B interaction, so FLARE is not activated. (**B**) Panel of β2AR agonists, partial agonists, and antagonist. Biased agonists preferentially recruit one downstream effector (such as β-arrestin2) over another. (**C**) β2AR-β-arrestin2 interaction timecourse with various ligands. HEK 293T cells expressing FLARE constructs were prepared as in Fig. 1D. 15 hours after transfection, 10 μM ligand was added at time = 0 minutes and remained on the cells for the duration of the experiment. The light window was 5 minutes, centered around the timepoint given on the x axis. 9 hours after initial addition of ligand, cells were mixed with luciferin substrate and analyzed for luciferase activity. Time courses are normalized with maximum signal ratio (SR) set to 1 and minimum SR set to 0. Each datapoint represents the mean of 4 replicates. Errors, STD. (**D**) Receptor-β-arrestin2 interaction time course with different receptors. Same as (C), except β2AR in the FLARE TF component was replaced with various other GPCRs, and ligand was added only briefly, from time = 0 to 5 minutes. (**E**) Same as (D), except ligand remained on the cells for the duration of the experiment (11 hours).

In addition to detecting the presence of ligands, scientists studying GPCRs are often interested in differentiating agonists from antagonists, and calculating EC50 and IC50 values. In Fig. 2E, we show that PPI-FLARE can be easily used to generate dose-response curves for the β2AR ligands isoproterenol (an agonist) and alprenolol (an antagonist).

In total, Fig. 2 extends PPI-FLARE to 12 different PPI pairs. Importantly, these results were all obtained simply by cloning genes into positions A and B of the constructs in Fig. 1A. We did not vary geometries or linkers or create panels of constructs for each PPI of interest. The success of all 12 PPI pairs we tested demonstrates that the PPI-FLARE scaffold is robust and highly modular in its design.

## PPI-FLARE for probing PPI dynamics

Because PPI-FLARE is gated by light, and light can be applied during any user-specified time window, we envisioned staggering the light window to read out different temporal regimes of a PPI time course. For instance, if an A-B PPI has the time course shown in Fig. 3A, left, a light window that overlaps with the peak of A-B interaction will result in FLARE activation and gene transcription, whereas a light window that is right-shifted to a later time period (Fig. 3A, right) will not. To test this concept, we used the β2AR-arrestin2 FLARE constructs, applied isoproterenol at t=0, and read out luciferase expression after shifting the light window to various time ranges (e.g., 0-5 minutes, 5-10 minutes, 15-20 minutes, etc.). Fig. 3C shows that the β2AR-β-arrestin2 interaction extent is high in the initial time period (0-5 minutes), and then falls off rapidly in the 5-10 minute range. This is consistent with previous reports using FRET^19^, translocation^20^, and proximity labeling^21^ assays, which show that the β2AR-arrestin interaction peaks between 1-3 minutes.

β2AR is a GPCR with multiple known ligands (Fig. 3B). Isoproterenol and isoetharine are both full agonists, while clenbuterol is a partial agonist, evoking a weaker downstream response and kinetically slower recruitment of arrestin^22^. Carvedilol is an inverse agonist, which can attenuate downstream signaling while still recruiting arrestin, albeit more weakly than a full agonist^23^. We used the light staggering scheme in Fig. 3A to examine the temporal response of the β2AR-arrestin2 interaction to these various ligands. As expected, the full agonists isoproterenol and isoetharine evoked strong responses (maximum ±ligand signal ratios ~18) and transient arrestin association. Clenbuterol induced a much smaller response and delayed onset of arrestin recruitment, consistent with its classification as a partial agonist. Interestingly, carvedilol produced a delayed but sustained interaction with β-arrestin2.

Different GPCRs are also thought to recruit arrestin with different kinetics. We performed the same FLARE time course assay, keeping the β-arrestin2-TEVp FLARE component constant, but varying the identity of the GPCR. Figure 3D shows that two Class A GPCRs, β2AR and DRD1, give strong FLARE responses within 5 minutes of ligand addition but much lower FLARE-driven luciferase expression within 5 minutes after ligand washout. This is consistent with previous studies showing that Class A GPCRs interact weakly and transiently with arrestin^24^. In contrast, the Class B GPCR AVPR2 recruits arrestin quickly and maintains a strong interaction with arrestin even 90 min after washout of the ligand vasopressin. The dopamine receptor DRD2 has not previously been classified as type A or B, but our data in Figure 3D strongly suggest that it is Class A, like DRD1, due to its transient association with arrestin. Figure 3E shows the same experiment, but with ligand left on the cells for the entire duration of the experiment. Now, FLARE-driven luciferase expression is observed even at 150 minutes for DRD1 and DRD2, but this is likely due to continual stimulation of fresh surface receptor pools with dopamine that remains in the culture media.

We conclude that, due to its time-gated transcriptional design, FLARE can resolve PPI dynamics, with a temporal resolution of 5 minutes, via a single timepoint readout that is simple and scalable (our assays in Fig. 3 were performed in 96-well plate format, with platereader readout). Excitingly, this feature enabled us to newly classify a receptor (DRD2) important in emotion, cognition, and neurological disease as a Class A-type GPCR.

## PPI-FLARE for genetic selections

Transcriptional assays such as yeast two-hybrid^25,26^ and the split-ubiquitin system^27,28^ have been adapted for PPI discovery via cell-based selections. In these experiments, each cell in a population expresses a constant “bait” protein and a variable “prey” protein. If in a particular cell, the prey interacts with bait, a transcription factor is reconstituted, and transcription of a survival gene such as HIS3 ensues. Though powerful, existing two-hybrid assays have two major limitations: high background, resulting in many false positive identifications, and no temporal resolution: is it impossible to distinguish between PPIs that occurred 1 minute after ligand addition versus 20 minutes after ligand addition, for example. PPI-FLARE offers a potential solution to these problems, because the light gate both reduces background/enhances specificity, *and* provides temporal precision, allowing the user to selectively focus on PPIs that occur during a 10-minute time window of their choosing.

We devised a preliminary “model selection” to see if PPI-FLARE could be combined with Fluorescence Activated Cell Sorting (FACS) to selectively enrich for cells expressing a matched PPI pair versus a mismatched, non-interacting protein pair (Fig. 4). We prepared HEK cells expressing the β2AR-β-arrestin2 FLARE components in Fig. 4A (matched PPI), and separate HEK cells expressing a mismatched β2AR-calmodulin FLARE pair, and mixed the two cell populations together in a 1:50 ratio (Fig. 4A). We then stimulated with isoproterenol and light for 10 minutes, and cultured for 9 additional hours to allow transcription and translation of the reporter gene, Citrine. Then cells were lifted and sorted on a FACS instrument to enrich cells with the highest Citrine expression. After a period of recovery and growth, we analyzed the cell population by qPCR to determine the ratio of arrestin cells (matched PPI FLARE) to calmodulin cells (mismatched PPI FLARE). Fig. 4C shows that a single round of FACS sorting can enrich the matched PPI cell population by 107-fold over the mismatched cell population. Such a large enrichment factor in a single round of selection suggests that it should be possible to use PPI FLARE to accurately discover PPIs that occur during selected time windows of interest.

**Figure 4:**
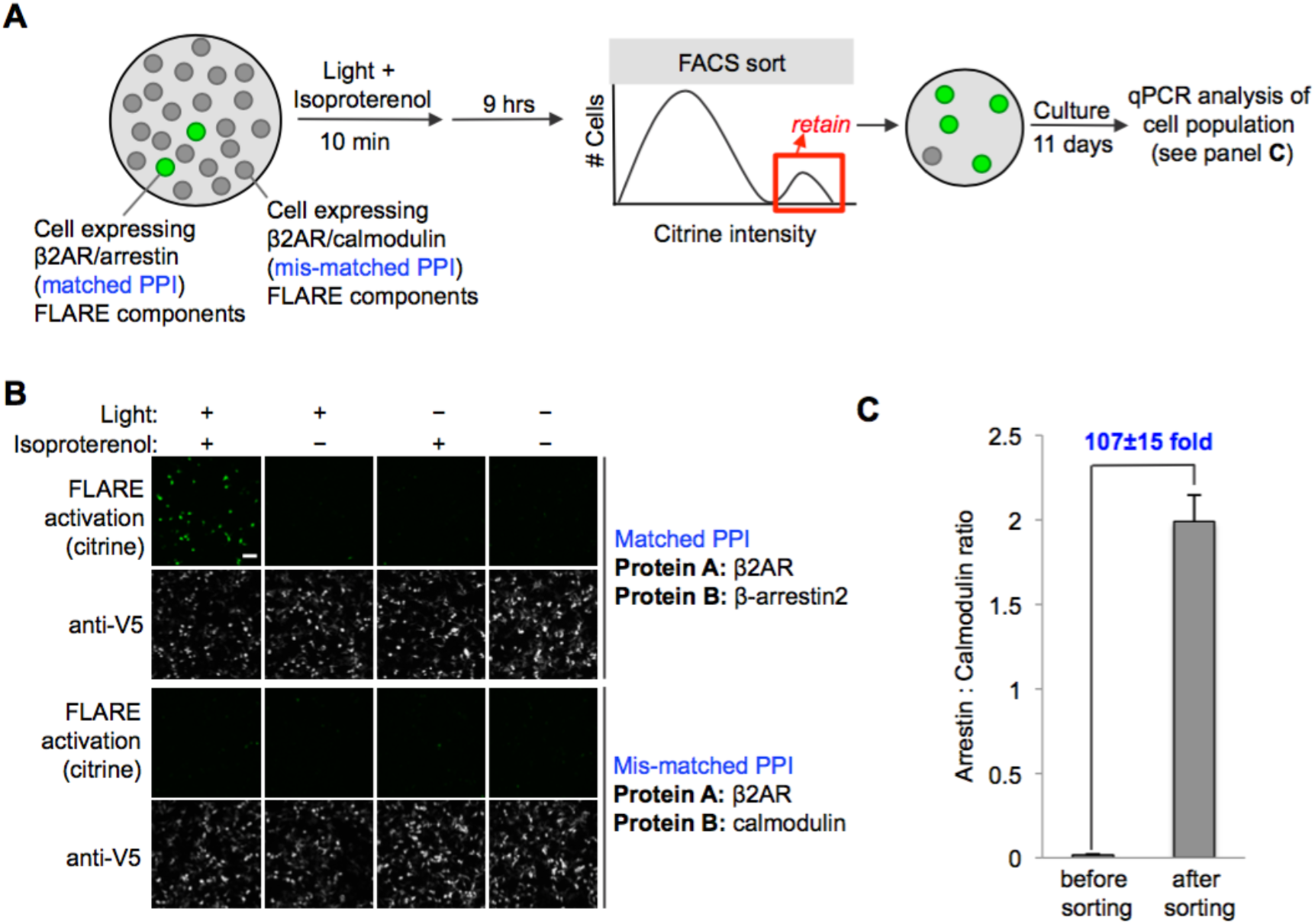
FLARE can be coupled to selections. (**A**) Selection scheme to enrich cells with PPI (protein-protein interaction) event during time window of interest over cells without PPI event. Culture dish containing mixed population of cells. Protein A (the “bait”) remains constant in the FLARE TF component, but protein B (“prey”) varies in the FLARE protease component. In our model selection, we used β-arrestin2 as the matched prey (interacts with β2AR), and calmodulin as the mis-matched prey (does not interact with β2AR). The culture dish is subjected to light and β2AR-stimulating ligand isoproterenol for 10 minutes, then cultured for an additional 9 hours. FACS sorting is performed to isolate cells with high Citrine expression, indicative of FLARE activation. After sorting, the cell population is expected to be strongly enriched in β-arrestin2-expressing cells over calmodulin-expressing cells. (**B**) Imaging of HEK 293T cells expressing FLARE components with matched bait + prey (top) versus mismatched bait + prey (bottom). Images were acquired 9 hours after stimulation with 10 µM isoproterenol and light (daylight lamp, 25W, 6500K, 480 nm/530 nm/590 nm). Anti-V5 antibody stains for the FLARE β2AR TF component. Scale bar, 100 µm. (**C**) qPCR analysis of mixed cell population before and after FACS sorting. Genomic DNA was extracted from cells and arrestin-TEVp: calmodulin-TEVp ratios were quantified by qPCR. Pre-sort ratio is 54:1; post-sort ratio is 2.0:1. Data represent the mean of three replicates. Error bar, STD.

## Comparison of PPI-FLARE to Tango and iTango

Tango is a transcriptional assay for reading out receptor activation that, like PPI-FLARE, uses PPI-driven proteolytic release of a transcription factor^9,11^ In contrast to PPI-FLARE, however, Tango lacks a light gate, and it utilizes a higher affinity TEVp-TEVcs pair (with K_m_ of 240 µM versus est. 450 µM for our system^14,29^) that reduces the dynamic range and increases the background of the assay. Furthermore, when applied to GPCRs, Tango requires a vasopressin receptor 2 tail domain (V2 tail) to be fused to the GPCR’s C-terminus, to artificially enhance arrestin recruitment and increase Tango signal. However, because Tango is used extensively in the GPCR field for its simple and scalable design^11^, we performed a side-by-side comparison to PPI FLARE. We prepared the β2AR-arrestin2 FLARE and Tango constructs shown in Supplementary Fig. 6A for the comparison.

HEK293T cells expressing FLARE or Tango constructs were treated with light and isoproterenol for 15 minutes, then cultured for 9 hours, before quantitation of luciferase expression (Supplementary Fig. 6B). The ±ligand signal ratio was 16.4-fold higher for FLARE than for Tango under these conditions. Since the Tango assay is typically performed with much longer periods of ligand stimulation^9,11^, we repeated the assay but increased the isoproterenol incubation to 18 hours (light period was still 15 minutes for FLARE). The Tango ±ligand signal ratio increased from 1.2 to 1.5, but was still far lower than the FLARE signal ratio (11.8). The predominant reason for this difference in performance appears to be high Tango background in the no-ligand condition. FLARE has much lower no-ligand luciferase expression because the eLOV domain protects the TEVcs from cleavage in the dark state. In contrast, Tango’s TEVcs is exposed throughout the entire period (hours to days) that the Tango components are expressed, leading to high background.

Recently, a light-gated version of Tango was reported, called iTango^13^. iTango was used to mark neurons exposed to dopamine in the mouse brain^13^. Though not explored for PPI detection in this study, the design of iTango is close enough to PPI-FLARE to suggest that it could potentially also be applied to PPI detection. Thus, we performed a side-by-side comparison of iTango and PPI-FLARE components, for detection of the isoproterenol-stimulated β2AR-arrestin interaction. The constructs we prepared for the comparison are shown in Supplementary Figure 6A, and the results shown in Supplementary Figure 6C. We found that under all conditions tested, iTango gave significantly higher background than PPI-FLARE. For example, there was 3 to 26-fold greater iTango-driven luciferase expression under dark/+ligand and light/no ligand conditions, than FLARE-driven luciferase expression under the same conditions. This resulted in 2 to 3-fold greater ±ligand signal ratios for PPI-FLARE, and 5 to 14-fold greater ±light signal ratios, compared to iTango (Supplementary Figure 6C).

The designs of iTango and PPI-FLARE differ in the following respects: iTango uses a split TEV protease with low catalytic turnover^13,30,31^, while FLARE uses a faster non-split TEV protease; iTango uses the V2 tail, like Tango, to artificially boost arrestin recruitment and increase signal, while PPI-FLARE does not require it; iTango uses iLID^13,15^ for light gating, while FLARE uses evolved eLOV^12^; and iTango uses a different TEVcs substrate (ENLYFQG versus ENLYFQM in FLARE). We hypothesize that the high background we observed with iTango derives mainly from its use of split TEV protease, whose fragments may have high affinity for one another, and reverse slowly, similar to other split proteins^32,33^. For example, in iTango’s no-ligand condition, a significant fraction of arrestin may nevertheless be complexed with β2AR, driven together by the split TEVp fragments to which they are fused. In the dark/+ligand condition, arrestin and β2AR may remain in complex long after isoproterenol is washed off, because split TEVp fragments probably dissociate slowly, giving TEVp a great deal of time to cleave proximal TEVcs, even in the dark state. Based on our collective observations, we conclude that PPI-FLARE is both more sensitive and more specific than Tango and iTango for PPI detection in cells.

## Discussion

The PPI-FLARE assay we have developed converts a specific, transient molecular interaction into a stable and amplifiable reporter gene signal. As such, PPI-FLARE is highly versatile, able to drive luciferase expression for platereader assays, or fluorescent protein expression for microscopy or FACS-based cell selections. Though not shown here, it should be facile to couple FLARE to the expression of other reporter genes as well, including APEX for EM^34,35^ and proteomics^36^, and various actuators such as opsins, toxins, and antibiotic resistance genes.

To develop PPI-FLARE, we optimized both the protease cleavage sequence and the light-sensitive LOV domain. The resulting tool is highly modular and generalizable; 12 of the 13 PPIs (including transmembrane as well as soluble proteins) we cloned into FLARE worked on the first try, without any optimization of geometries or linkers. PPI-FLARE is highly specific for direct PPIs, and is not activated merely by protein proximity (e.g., colocalization to the plasma membrane – Fig. 1F). We showed that only 5 minutes of light coinciding with a PPI event are sufficient to produce a stable signal that lasts for 9 hours to days.

PPI-FLARE produces larger signal-to-noise ratios than other transcriptional PPI assays. When Presto-Tango was applied to 167 different GPCRs, for example, the majority of them gave signal ratios between 1.3 and 5 (Ref. 11). In contrast, PPI-FLARE gave signal ratios >15 for five of the eight GPCRs that we randomly selected, and signal ratios >10 for other PPI pairs (Figs. 2B and C). The greater signal ratios result from two factors. First, noise is lower for PPI-FLARE because the LOV domain prevents proteolytic cleavage except during the brief light window. Second, signal is higher for PPI-FLARE because our TEVp-TEVcs pair has higher catalytic turnover (est. 10.8 min^-1^ for PPI-FLARE compared to 0.84 min^-1^ for Tango; iTango uses split-TEVp^13^ which is expected to have even slower turnover). As a result of improved signal ratio, PPI-FLARE can be used without artificial enhancement of native affinity (e.g., V2 tail), in contrast to both Tango and iTango. And due to its higher sensitivity, PPI-FLARE can detect the activation of receptors that are opaque to other methods, such as ADRA1A and AGTR2 in Fig. 2D, which gave no ±ligand signal difference with Presto-Tango^11^.

Protein Complementation Assays (PCAs) are a commonly used alternative assay to detect and study cellular PPIs. Irreversible PCAs such as bimolecular fluorescence complementation (BiFC)^33^ and split HRP^32^ also integrate molecular events and produce stable signals, but unlike PPI-FLARE, they trap PPI partners in an irreversible complex that can perturb cellular signaling and give rise to false positives. Reversible PCAs such as split luciferase^37^ do not trap, but they also do not produce stable signals that enable single time point readout as PPI-FLARE does. By integrating through a transcription factor, PPI-FLARE is unique in combining a non-trapping design with stable, integrated signal generation.

PPI dynamics are central to signal transduction but previous transcriptional assays were unable to probe this aspect of their biology. Due to PPI-FLARE’s light gate, we showed that it is possible to “time stamp” PPI events, and even to reconstruct PPI time courses. We also made the interesting observation that just a simple four-condition, single timepoint experiment (±ligand and ±light) could reveal information about PPI dynamics: a transient PPI gives a small ±ligand (dark state) signal ratio, whereas a sustained PPI produces a larger ratio, presumably because imperfect LOV caging gives rise to some proteolytic cleavage when PPIs remain in complex for long periods of time.

Future applications of PPI-FLARE might include examination of GPCR “biased” signaling^38^. Here, we used only arrestin-TEVp with our GPCR panel, but it should be possible to replace the arrestin gene with Gi, Gs, or Gq and compare intensities and timecourses of G protein versus arrestin recruitment to GPCRs in response to various ligands. Our model selection using FACS also suggests that PPI-FLARE will have utility for PPI discovery and other functional genomics applications. Notably, while yeast two hybrid and split ubiquitin assays have been employed extensively for yeast-based selections^27,39^, we are not aware of any applications of these platforms or related ones to mammalian cell-based selections, perhaps because the signal-to-noise ratios of these assays are not sufficiently high.

Despite its advantages, PPI-FLARE has also notable limitations. First, it is a recombinant assay, and PPI partners are fused to rather large engineered proteins. Artifacts could arise from overexpression or steric interference with PPI trafficking or interactions. Second, we showed that PPI-FLARE could be applied to both transmembrane and soluble proteins, but the latter require artificial sequestration onto a membrane, to prevent nuclear translocation in the basal state. Third, due to its intermolecular design, PPI-FLARE signal is expression-level dependent. We found that stable expression of tool components improves assay performance due to lowering of the background. Fourth, a 5-minute time gate permits study of certain dynamic processes, but many cellular events occur on a faster timescale. Engineering of TEVp to increase its catalytic turnover rate should yield a future PPI-FLARE tool with even shorter tagging times.

## Acknowledgements

We thank K. K. Kumar and B. Kobilka for helpful discussions. FACS experiments were performed in the Stanford Shared FACS Facility. Rat β-arrestin2 and β2AR genes were a gift from R. Lefkowitz, Duke University. The EGFR gene was a gift from M. Meyerson, Dana-Farber Cancer Institute. The Grb2 gene was a gift from B. Mayer, University of Connecticut. The luciferase gene was a gift from M. Sato, University of Tokyo. DRD1, DRD2, NMBR, MLNR, AVPR2, AGTR2, and ADRA1A genes were a gift from B. Roth, University of North Carolina. A.Y.T. received funding from Stanford University. A.Y.T. is a Chan Zuckerberg Biohub investigator.

## Author Contributions

M.W.K. and W.W. performed all experiments except those noted. M.I.S. cloned constructs for the LOV domain comparison, and performed FACS sorting. R.C. performed the mitochondria FKBP-FRB FLARE experiment. M.V.Z. provided advice and guidance. M.W.K., W.W. and A.Y.T. designed the research, analyzed the data, and wrote the paper. All authors edited the paper.

## Supplementary Figures

**Supplementary figure 1:**
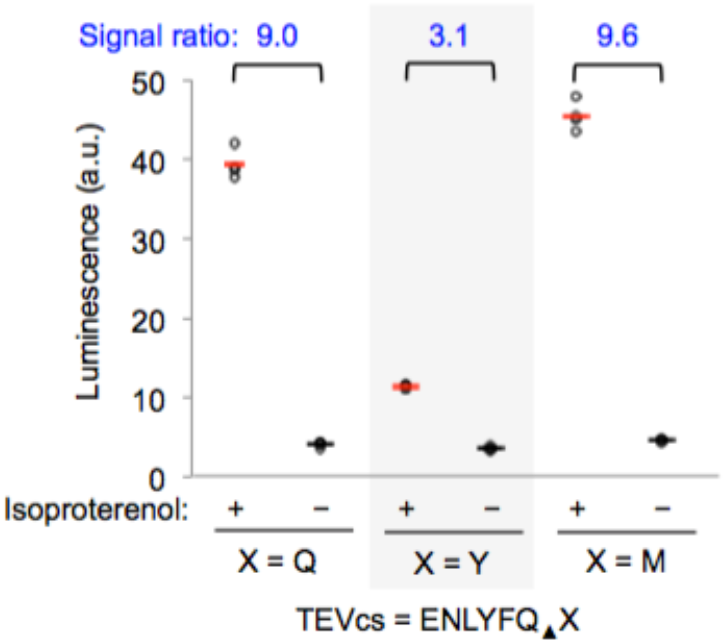
Testing alternative TEVcs sequences. Three alternative TEVcs sequences that differ at the P1´ site were tested in the context of β2AR-β-arrestin2 FLARE. HEK cells were prepared as in Fig. 1D and stimulated with 10 µM isoproterenol and blue LED light for 5 minutes. Nine hours later, cells were analyzed for luciferase activity. Each condition was replicated four times. We then used the TEVcs sequence with X=M for all experiments in this study, except where indicated.

**Supplementary figure 2:**
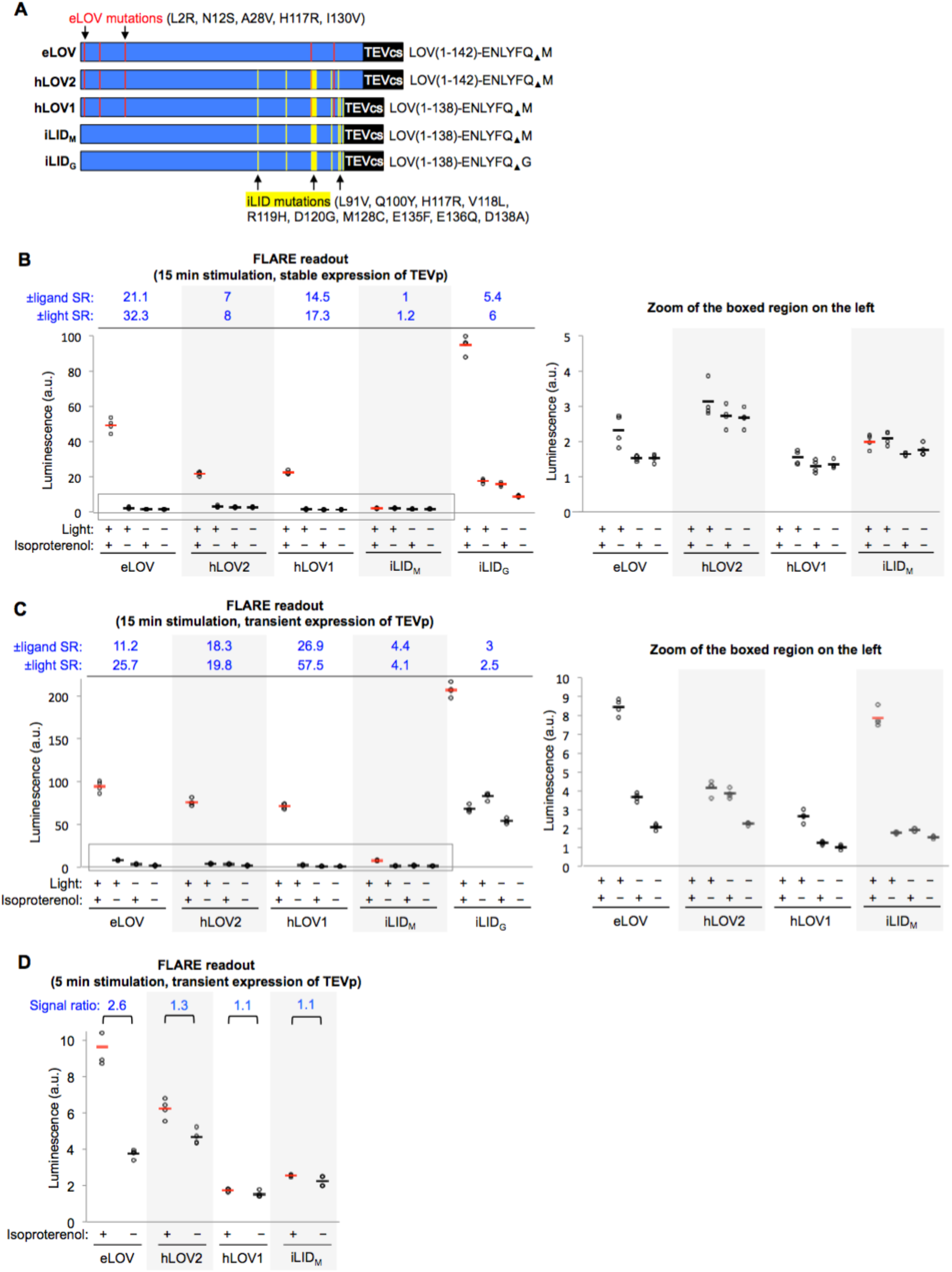
Testing alternative LOV domains. (**A**) Five LOV-TEVcs fusions compared. eLOV (top) was engineered by directed evolution in a previous study^12^, and is used in all FLARE experiments in this study, except where indicated. The red lines indicate where the eLOV sequence differs from that of AsLOV2(G126A/N136E)^43^, the template used for directed evolution. iTANGO^13^ uses the LOV domain from iLID^15^(bottom two constructs) and its TEVcs “bites back” 6 amino acids into LOV’s Jα helix. Yellow lines indicate where iLID’s LOV sequence differs from that of AsLOV2. hLOV1 and hLOV2 are two hybrid LOV domains that merge the features of eLOV and iLID. TEVcs is the same in the top four constructs but has Gly instead of Met in the P1´ position in the bottom construct. (**B**) Comparison of five LOV-TEVcs fusions, with luciferase readout, and stable/low expression of β-arrestin2-TEVp. The boxed region is enlarged on the right. HEK 293T cells were prepared as in Fig. 1D, with β-arrestin2-TEVp stably expressed and FLARE β2AR-TF (containing one of five LOV-TEVcs sequences from (A)) and UAS-luciferase transiently expressed. 18 hours post-transfection, cells were stimulated with 15 minutes of isoproterenol and ambient light. Nine hours later, cells were analyzed for luciferase activity. Each condition was replicated four times. ± ligand signal ratios (SR) and ± light signal ratios for each construct quantified across top. (**C**) Same as (B), but with transient overexpression of β-arrestin2-TEVp component, instead of stable/low expression. The boxed region is enlarged on the right. (**D**) Same as (C) but with 5 min instead of 15 min light stimulation.

**Supplementary figure 3:**
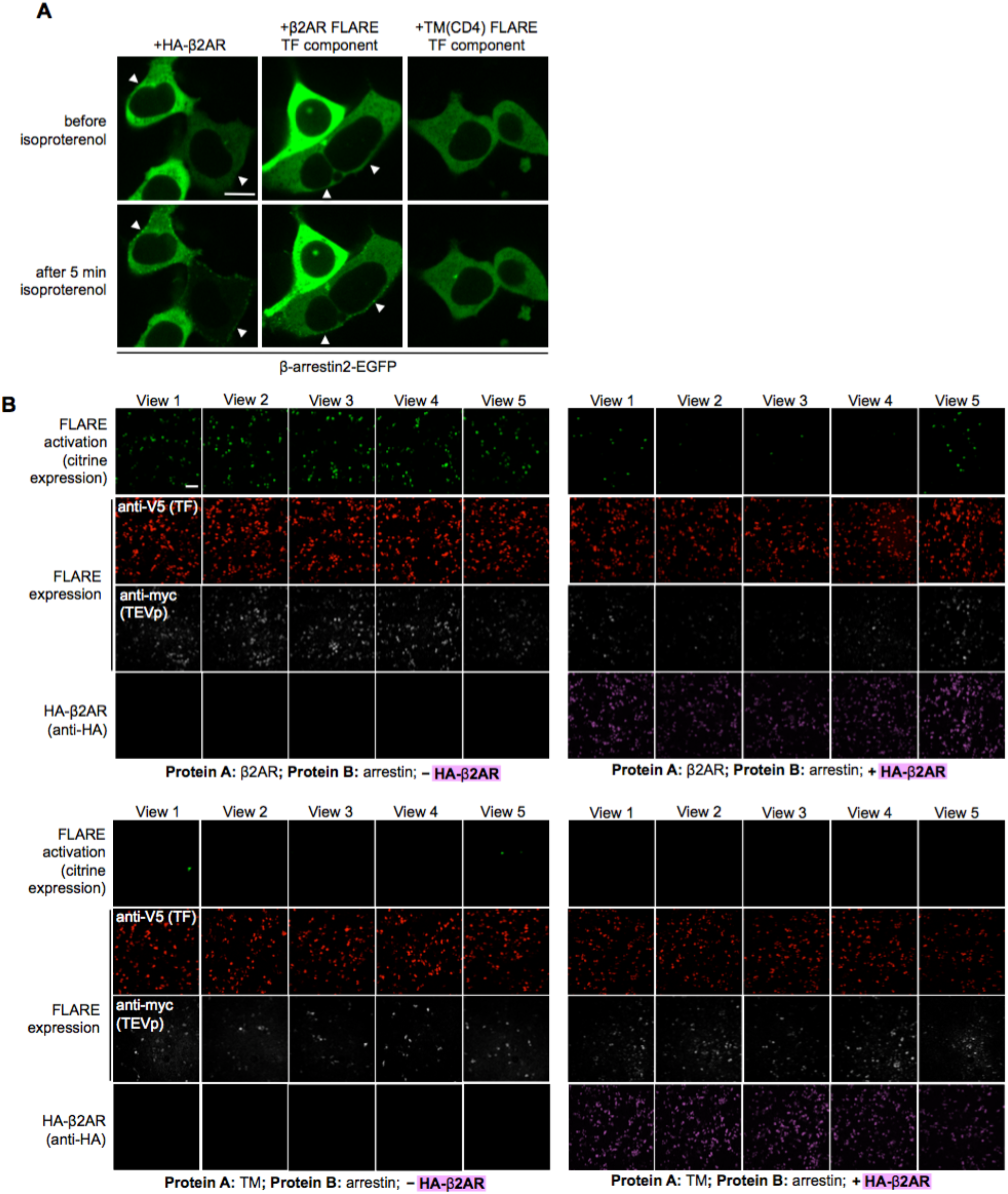
Additional data related to Fig. 1F. (**A**) HA-β2AR construct recruits β-arrestin2 -EGFP to the plasma membrane. GFP images of HEK 293T cells transiently expressing rat β-arrestin2-EGFP along with one of the following: HA-β2AR, β2AR FLARE TF component (from Fig. 1B), or TM FLARE TF component (TM from CD4, used in Fig. 1F). Live cell GFP images were acquired before and after incubation with 10 µM isoproterenol to activate β2AR. Arrowheads point to regions showing re-localization of β-arrestin2-GFP. Scale bar, 10 µm. (**B**) Additional fields of view for the experiment shown in Fig. 1F. Scale bar, 100 µm.

**Supplementary figure 4:**
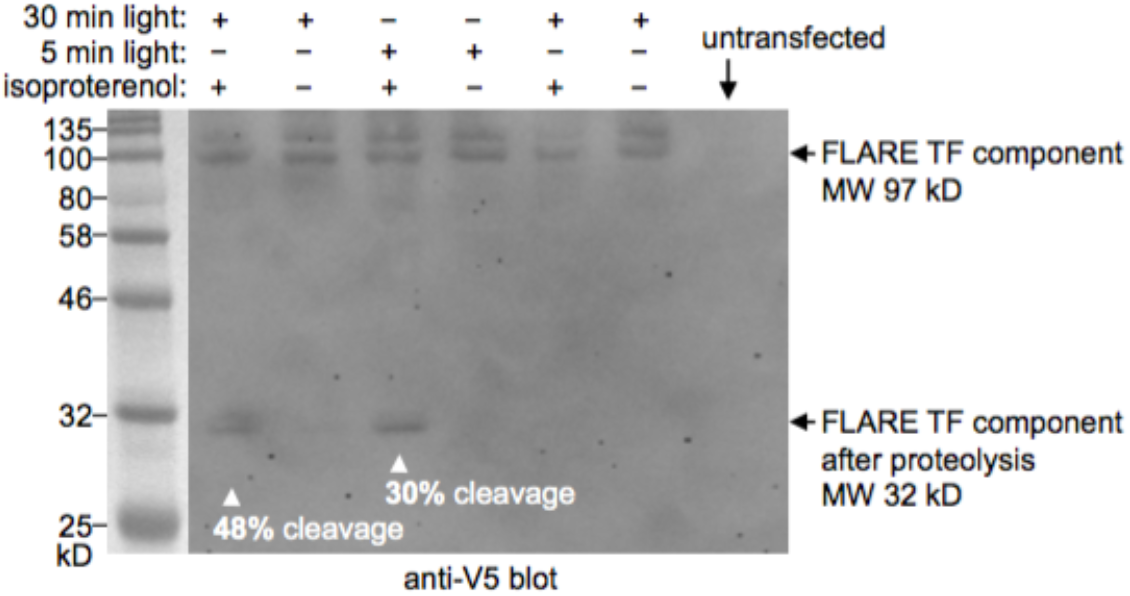
Western blot quantification of cleavage extent. HEK 293T cells were transiently transfected (using PEI max) with the PPI-FLARE constructs shown in Fig. 1B. 18 hrs post-transfection, cells were stimulated with 10 µM isoproterenol and blue light (467 nm, 60 mW/cm^2^, 10% duty cycle) for 5 or 30 minutes total. Cells were then immediately lysed in the presence of 20 mM iodoacetamide TEVp inhibitor and run on 8% SDS-PAGE. Anti-V5 blot visualizes the FLARE TF component, which is 97 kD before cleavage and 32 kD after cleavage at the TEVcs. Negative controls omit isoproterenol or light.

**Supplementary figure 5:**
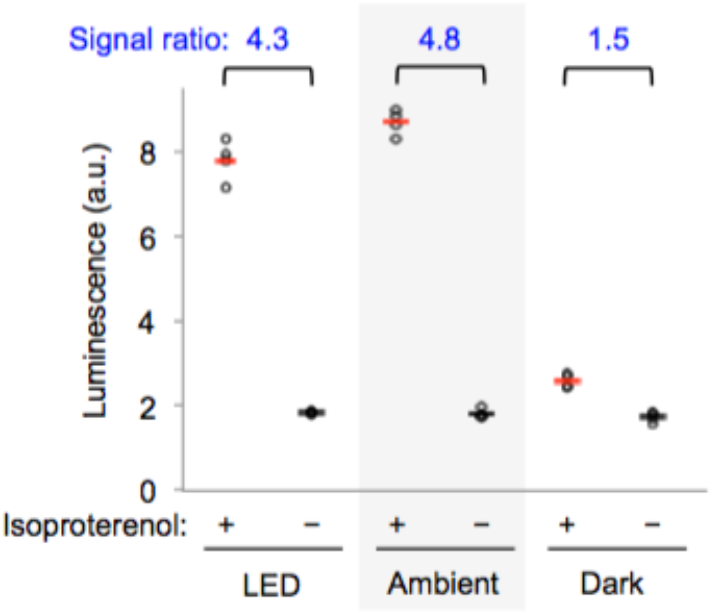
Ambient room light activates FLARE. HEK 293T cells were prepared as in Fig. 1D. 15 hours post-transfection, cells were stimulated with 5 minutes of either ambient room light or blue LED light (467 nm, 60 mW/cm^2^, 10% duty cycle) concurrently with 10 µM isoproterenol. Nine hours later, cells were analyzed for luciferase activity. Each condition was replicated four times.

**Supplementary figure 6:**
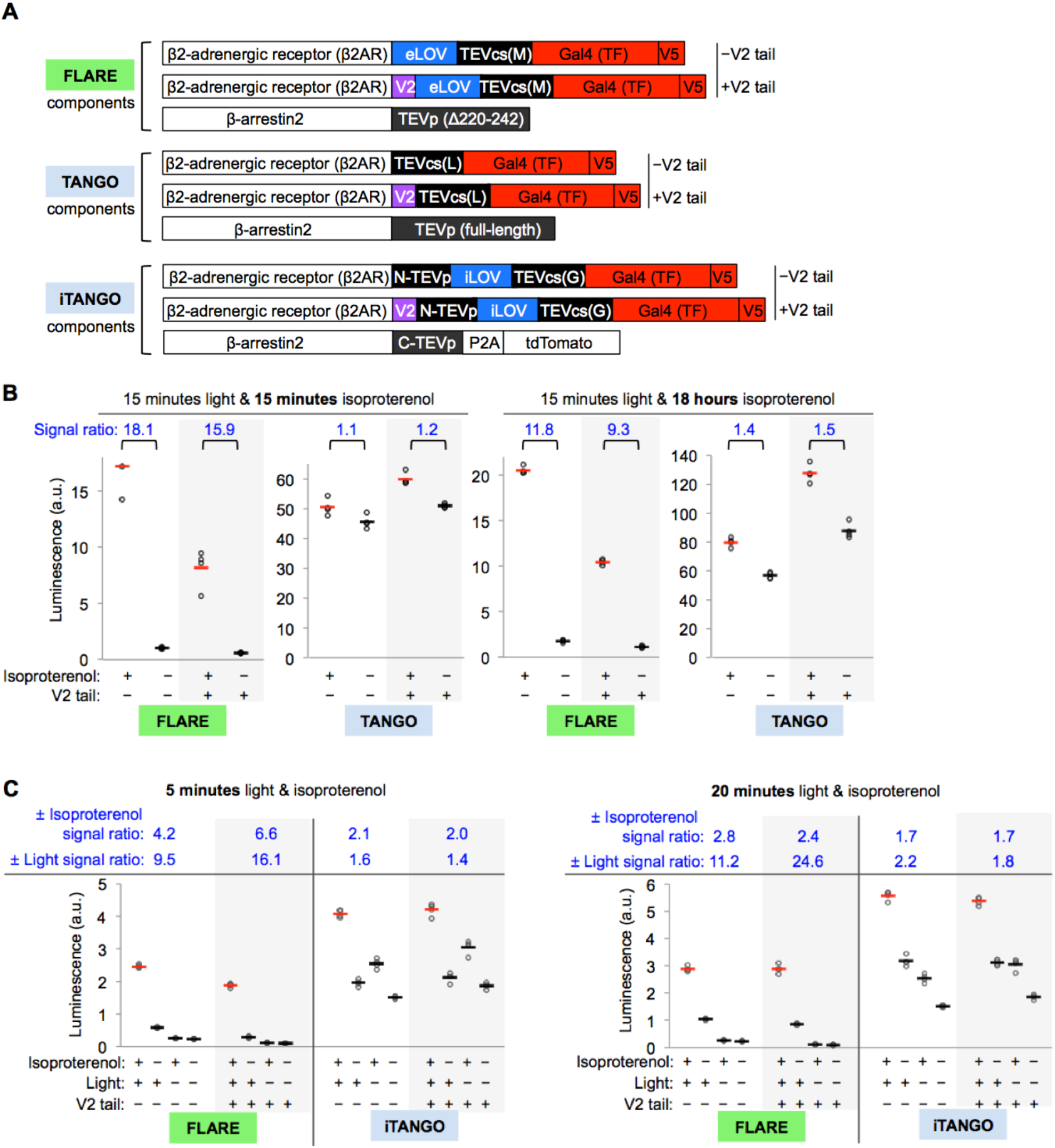
PPI-FLARE comparison to TANGO and iTango. (**A**) FLARE, TANGO, and iTANGO constructs used to detect β2AR-β-arrestin2 interaction. The β2AR fusions were each prepared with and without the vasopressin receptor tail (V2, purple) that enhances arrestin recruitment^11^. These FLARE, TANGO, and iTANGO constructs differ only in their TEVcs, TEVp, and LOV sequences; arrestin, β2AR, and TF domains are constant. In comparison to FLARE, TANGO uses full-length TEVp and a lower-affinity TEVcs with Leu instead of Met at the P1´ site. TANGO has no light gating. In comparison to FLARE, iTango uses a split TEVp, a higher-affinity TEVcs with Gly at the P1´ site, and the LOV sequence from iLID (iLOV)^15^. (**B**) FLARE versus TANGO comparison. HEK 293T cells stably expressing the protease component of FLARE or TANGO were transiently transfected with the corresponding TF component and UAS-luciferase. 18 hours post-transfection, cells were stimulated with 15 minutes of light (daylight lamp) and isoproterenol, then analyzed for luciferase activity 9 hours later (left). Alternatively (right), cells were stimulated with 15 minutes of light in the presence of isoproterenol, and isoproterenol remained on the cells for another 18 hours, before luciferase detection (to match published conditions for TANGO^9,10^). Each condition was replicated four times. ± isoproterenol signal ratios are quantified at top. (**C**) FLARE versus iTANGO comparison. Constructs shown in (A) were introduced by lipofectamine transfection into HEK 293T cells along with UAS-luciferase. 18 hrs post-transfection, cells were stimulated with either 5 minutes (left) or 20 minutes (right) of isoproterenol and light (daylight lamp). Nine hours later, cells were analyzed for luciferase activity. Each condition was replicated four times. ±Isoproterenol and ±light signal ratios are quantified at top.

## References

(1) Truong, K., and Ikura, M. (2001) The use of FRET imaging microscopy to detect protein-protein interactions and protein conformational changes in vivo. Curr. Opin. Struct. Biol.

(2) Pfleger, K. D. G., and Eidne, K. A. (2006) Illuminating insights into protein-protein interactions using bioluminescence resonance energy transfer (BRET). Nat. Methods 3, 165–174.

(3) Krichevsky, O., and Bonnet, G. (2002) Fluorescence correlation spectroscopy: the technique and its applications. Rep. Prog. Phys 65, 251–297.

(4) Shekhawat, S. S., and Ghosh, I. (2011) Split-protein systems: Beyond binary protein-protein interactions. Curr. Opin. Chem. Biol.

(5) Teruel, M. N., and Meyer, T. (2000) Translocation and reversible localization of signaling proteins: a dynamic future for signal transduction. Cell 103, 181–4.

(6) Yang, X., Jost, A. P.-T., Weiner, O. D., and Tang, C. (2013) A light-inducible organelle-targeting system for dynamically activating and inactivating signaling in budding yeast. Mol. Biol. Cell 24, 2419–2430.

(7) Miller, J., and Stagljar, I. (2004) Using the Yeast Two-Hybrid System to Identify Interacting Proteins. Protein-Protein Interact. 261, 247–262.

(8) Petschnigg, J., Groisman, B., Kotlyar, M., Taipale, M., Zheng, Y., Kurat, C. F., Sayad, A., Sierra, J. R., Usaj, M. M., Snider, J., Nachman, A., Krykbaeva, I., Tsao, M.-S., Moffat, J., Pawson, T., Lindquist, S., Jurisica, I., and Stagljar, I. (2014) The mammalian-membrane two-hybrid assay (MaMTH) for probing membrane-protein interactions in human cells. Nat. Methods 11, 585–92.

(9) Barnea, G., Strapps, W., Herrada, G., Berman, Y., Ong, J., Kloss, B., Axel, R., and Lee, K. J. (2008) The genetic design of signaling cascades to record receptor activation. Proc. Natl. Acad. Sci. U. S. A. 105, 64–9.

(10) Inagaki, H. K., Ben-Tabou De-Leon, S., Wong, A. M., Jagadish, S., Ishimoto, H., Barnea, G., Kitamoto, T., Axel, R., and Anderson, D. J. (2012) Visualizing neuromodulation in vivo: TANGO-mapping of dopamine signaling reveals appetite control of sugar sensing. Cell 148, 583–595.

(11) Kroeze, W. K., Sassano, M. F., Huang, X.-P., Lansu, K., McCorvy, J. D., Giguère, P. M., Sciaky, N., and Roth, B. L. (2015) PRESTO-Tango as an open-source resource for interrogation of the druggable human GPCRome. Nat. Struct. Mol. Biol.

(12) Wang, W., Wildes, C. P., Pattarabanjird, T., Sanchez, M. I., Glober, G. F., Matthews, G. A., Tye, K. M., and Ting, A. Y. (2017) A light- and calcium-gated transcription factor for imaging and manipulating activated neurons. Nat. Biotechnol.

(13) Lee, D., Creed, M., Jung, K., Stefanelli, T., Wendler, D. J., Oh, W. C., Mignocchi, N. L., Lüscher, C., and Kwon, H.-B. (2017) Temporally precise labeling and control of neuromodulatory circuits in the mammalian brain. Nat. Methods 14, 495–503.

(14) Kapust, R. B., Tözsér, J., Copeland, T. D., and Waugh, D. S. (2002) The P1′ specificity of tobacco etch virus protease. Biochem. Biophys. Res. Commun. 294, 949–955.

(15) Guntas, G., Hallett, R. A., Zimmerman, S. P., Williams, T., Yumerefendi, H., Bear, J. E., and Kuhlman, B. (2015) Engineering an improved light-induced dimer (iLID) for controlling the localization and activity of signaling proteins. Proc. Natl. Acad. Sci. 112, 112–117.

(16) Pudasaini, A., and Zoltowski, B. D. (2013) Zeitlupe senses blue-light fluence to mediate circadian timing in arabidopsis thaliana. Biochemistry 52, 7150–7158.

(17) Hosoi, H., Dilling, M. B., Shikata, T., Liu, L. N., Shu, L., Ashmun, R. A., Germain, G. S., Abraham, R. T., and Houghton, P. J. (1999) Rapamycin causes poorly reversible inhibition of mTOR and induces p53- independent apoptosis in human rhabdomyosarcoma cells. Cancer Res. 59, 886–894.

(18) Shenoy, S. K., and Lefkowitz, R. J. (2011) β-arrestin-mediated receptor trafficking and signal transduction. Trends Pharmacol. Sci.

(19) Reiner, S., Ambrosio, M., Hoffmann, C., and Lohse, M. J. (2010) Differential signaling of the endogenous agonists at the beta2-adrenergic receptor. J. Biol. Chem. 285, 36188–36198.

(20) Eichel, K., Jullié, D., and von Zastrow, M. (2016) β-Arrestin drives MAP kinase signalling from clathrin-coated structures after GPCR dissociation. Nat. Cell Biol. 18, 303–310.

(21) Lobingier, B. T., Hü, R., Eichel, K., Miller, K. B., Ting, A. Y., Von Zastrow, M., and Krogan, N. J. (2017) An Approach to Spatiotemporally Resolve Protein Interaction Networks in Living Cells Resource An Approach to Spatiotemporally Resolve Protein Interaction Networks in Living Cells. Cell 169, 350–360.

(22) Kaya, A. I., Onaran, H. O., Özcan, G., Ambrosio, C., Costa, T., Balli, S., and Uǧur, Ö. (2012) Cell contact-dependent functional selectivity of β2-adrenergic receptor ligands in stimulating cAMP accumulation and extracellular signal-regulated kinase phosphorylation. J. Biol. Chem. 287, 6362–74.

(23) Wisler, J. W., DeWire, S. M., Whalen, E. J., Violin, J. D., Drake, M. T., Ahn, S., Shenoy, S. K., and Lefkowitz, R. J. (2007) A unique mechanism of β-blocker action: carvedilol stimulates β-arrestin signaling. Proc. Natl. Acad. Sci. U. S. A. 104, 16657–16662.

(24) Oakley, R. H., Laporte, S. A., Holt, J. A., Caron, M. G., and Barak, L. S. (2000) Differential affinities of visual arrestin, βarrestin1, and βarrestin2 for G protein-coupled receptors delineate two major classes of receptors. J. Biol. Chem. 275, 17201–17210.

(25) Fields, S., and Song, O. (1989) A novel genetic system to detect protein-protein interactions. Nature 340, 245–246.

(26) Luo, Y., Batalao, A., Zhou, H., and Zhu, L. (1997) Mammalian two-hybrid system: A complementary approach to the yeast two- hybrid system. Biotechniques 22, 350–352.

(27) Fetchko, M., and Stagljar, I. (2004) Application of the split-ubiquitin membrane yeast two-hybrid system to investigate membrane protein interactions. Methods 32, 349–362.

(28) Petschnigg, J., Groisman, B., Kotlyar, M., Taipale, M., Zheng, Y., Kurat, C. F., Sayad, A., Sierra, J. R., Usaj, M. M., Snider, J., Nachman, A., Krykbaeva, I., Tsao, M.-S., Moffat, J., Pawson, T., Lindquist, S., Jurisica, I., and Stagljar, I. (2014) The mammalian-membrane two-hybrid assay (MaMTH) for probing membrane-protein interactions in human cells. Nat. Methods 11, 585–592.

(29) Kapust, R. B., Tözsér, J., Fox, J. D., Anderson, D. E., Cherry, S., Copeland, T. D., and Waugh, D. S. (2001) Tobacco etch virus protease: mechanism of autolysis and rational design of stable mutants with wild-type catalytic proficiency. Protein Eng. 14, 993–1000.

(30) Scheek, S., Wehr, M. C., Laage, R., Bolz, U., Fischer, T. M., Gru, S., Bach, A., Nave, K., and Rossner, M. J. (2006) Monitoring regulated protein-protein interactions using split TEV. Nat. Methods 3.

(31) Gray, D. C., Mahrus, S., and Wells, J. A. (2010) Activation of specific apoptotic caspases with an engineered small-molecule-activated protease. Cell 142, 637–646.

(32) Martell, J. D., Yamagata, M., Deerinck, T. J., Phan, S., Kwa, C. G., Ellisman, M. H., Sanes, J. R., and Ting, A. Y. (2016) A split horseradish peroxidase for the detection of intercellular protein–protein interactions and sensitive visualization of synapses. Nat. Biotechnol. 32, 13819–13840.

(33) Kerppola, T. K. (2008) Bimolecular fluorescence complementation (BiFC) analysis as a probe of protein interactions in living cells. Annu Rev Biophys 37, 465–87.

(34) Martell, J. D., Deerinck, T. J., Sancak, Y., Poulos, T. L., Mootha, V. K., Sosinsky, G. E., Ellisman, M. H., and Ting, A. Y. (2012) Engineered ascorbate peroxidase as a genetically encoded reporter for electron microscopy. Nat. Biotechnol. 30, 1143–8.

(35) Lam, S. S., Martell, J. D., Kamer, K. J., Deerinck, T. J., Ellisman, M. H., Mootha, V. K., and Ting, A. Y. (2014) Directed evolution of APEX2 for electron microscopy and proximity labeling. Nat. Methods 12, 51–54.

(36) Rhee, H., Zou, P., Udeshi, N. D., Martell, J. D., Mootha, V. K., Carr, S. A., and Ting, A. Y. (2013) Proteomic mapping of mitochondria in living cells via spatially restricted enzymatic tagging. Science 339, 1328–31.

(37) Ozawa, T., Kaihara, A., Sato, M., Tachihara, K., and Umezawa, Y. (2001) Split luciferase as an optical probe for detecting protein-protein interactions in mammalian cells based on protein splicing. Anal Chem 73, 2516–2521.

(38) Bologna, Z., Teoh, J.-P., Bayoumi, A. S., Tang, Y., and Kim, I.-M. (2017) Biased G Protein-Coupled Receptor Signaling: New Player in Modulating Physiology and Pathology. Biomol. Ther. (Seoul). 25, 12–25.

(39) Ito, T., Chiba, T., Ozawa, R., Yoshida, M., Hattori, M., and Sakaki, Y. (2001) A comprehensive two-hybrid analysis to explore the yeast protein interactome. Proc. Natl. Acad. Sci. 98, 4569–4574.

(40) Kennedy, M. J., Hughes, R. M., Peteya, L. A., Schwartz, J. W., Ehlers, D., and Tucker, C. L. (2011) Rapid blue light induction of protein interactions in living cells. Nat. Methods 7, 973–975.

(41) Fisher, G. W., Adler, S. A., Fuhrman, M. H., Waggoner, A. S., Bruchez, M. P., and Jarvik, J. W. (2010) Detection and quantification of beta2AR internalization in living cells using FAP-based biosensor technology. J. Biomol. Screen. 15, 703–709.

(42) Gether, U., Lin, S., and Kobilka, B. K. (1995) Fluorescent labeling of purified beta 2 adrenergic receptor. Evidence for ligand-specific conformational changes. J. Biol. Chem. 270, 28268–28275.

(43) Strickland, D., Yao, X., Gawlak, G., Rosen, M. K., Gardner, K. H., and Sosnick, T. R. (2010) Rationally improving LOV domain-based photoswitches. Nat. Methods 7, 623–6.

